# Mantis shrimp navigate home using celestial and idiothetic path integration

**DOI:** 10.1101/2020.03.06.981043

**Authors:** Rickesh N. Patel, Thomas W. Cronin

## Abstract

Path integration is a robust mechanism that many animals employ to return to specific locations, typically their homes, during navigation. This efficient navigational strategy has never been demonstrated in a fully aquatic animal, where sensory cues used for orientation may differ dramatically from those available above the water’s surface. Here we report that the mantis shrimp, *Neogonodactylus oerstedii*, uses path integration informed by a hierarchical reliance on the sun, overhead polarization patterns, and idiothetic (internal) orientation cues to return home when foraging, making them the first fully aquatic path-integrating animals yet discovered. We show that mantis shrimp rely on navigational strategies closely resembling those used by insect navigators, opening a new avenue for the investigation of the neural basis of navigation behaviors and the evolution of these strategies in arthropods and potentially other animals as well.

## Introduction

Many central place foragers, animals that to return to a home location between foraging bouts, efficiently navigate to their homes using path integration. During path integration, an animal monitors its angular and linear movements using compass and odometer cues. From this information, a home vector, the most direct path back to the reference point, is calculated and continually updated, allowing the animal to return to its original location [1-4].

Path integration has been most thoroughly studied in social hymenopterans, which primarily rely on celestial cues for orientation and on idiothetic cues for odometry [2, 5-8]. This navigational strategy is also used by other terrestrial taxa [1, 3, 4], but has not been demonstrated in any fully aquatic animal. Sensory cues underwater differ dramatically from those available in terrestrial environments; for example, visual cues which are prominent on land are obscured over relatively short distances underwater.

Stomatopods, better known as mantis shrimp, are predatory crustaceans that mostly inhabit shallow marine waters. Many stomatopod species occupy small holes in their benthic environments where they are safely concealed from their predators. Most species leave these burrows, risking predation, for tasks such as foraging and finding mates [9-11]. These trips away from the burrow may extend to four meters or more in some *Neogonodactylus* species, a substantial distance for animals typically around three to five centimeters long [9-11]. We found that *Neogonodactylus oerstedii* in shallow waters off the Florida Keys make multiple excursions from and back to a home burrow and that these excursions extend up to a few meters from the burrow (Fig. 1). Further, the densities of occupied burrows were fairly high in some locations, with burrows of multiple animals as close as 10 centimeters to one another. Due to the aggressive territoriality *Neogonodactylus* (and many other stomatopods) exhibit when defending their burrows and the powerful weaponry they possess [12], accurate navigation back to the correct home burrow is an important task for a mantis shrimp.

**Figure 1.**
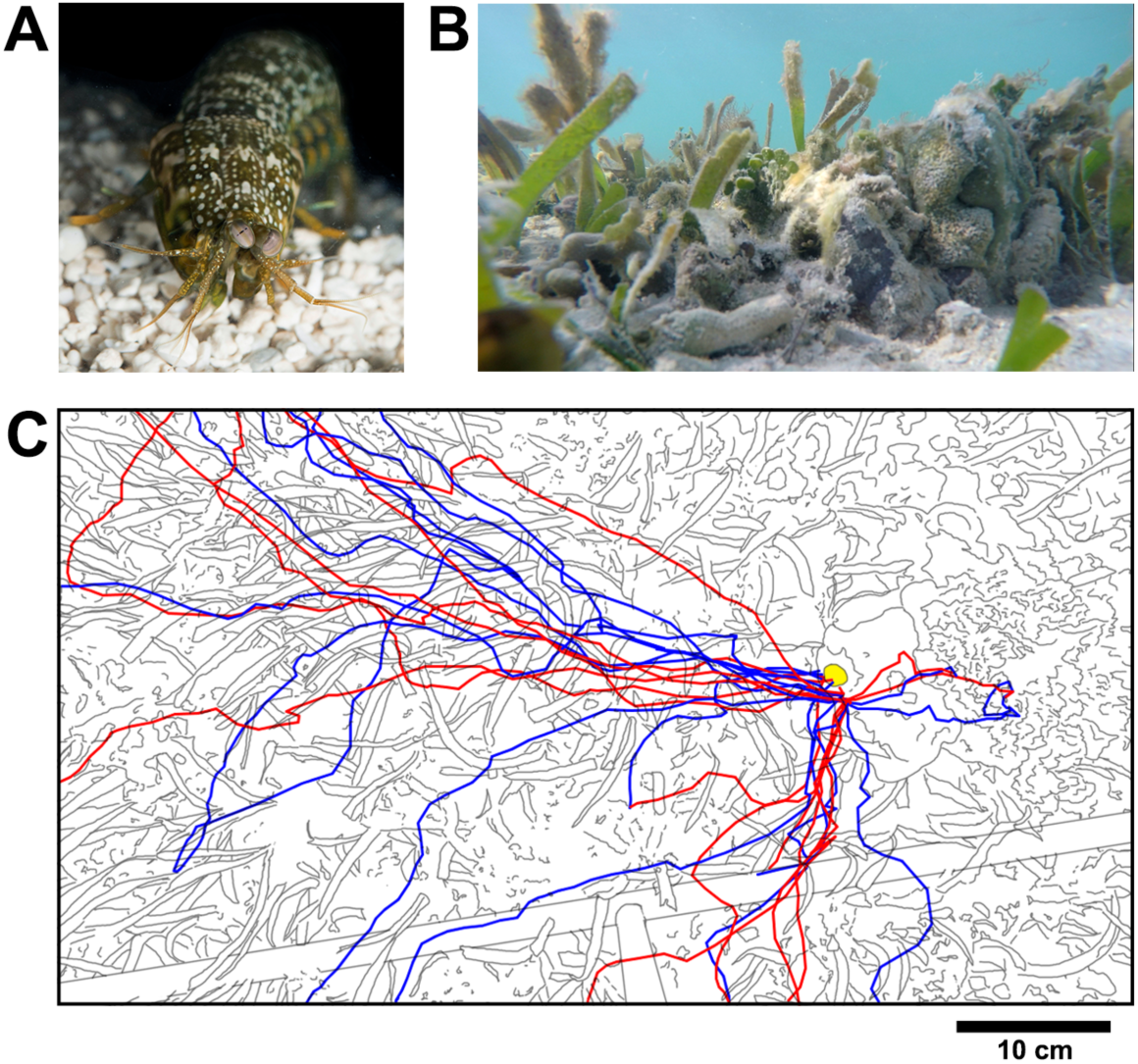
*Neogonodactylus oerstedii* makes multiple foraging trips from and to its burrow in nature. **(A)** *Neogonodactylus oerstedii* **(B)** *N. oerstedii* peering out of its burrow in nature. **(C)** Foraging routes of *N. oerstedii* from (blue tracings) and to (red tracings) its burrow (filled in yellow) in nature viewed from above. Data were obtained over three hours during a single evening.

These considerations led us to investigate the mechanisms used by *Neogonodactylus oerstedii* to navigate back to its home when foraging.

## Results and Discussion

### Mantis shrimp navigate using path integration

In initial experiments, we placed individual *N. oerstedii* in relatively featureless sandy-bottomed circular arenas filled with sea water in a glass-roofed greenhouse. Vertical burrows were buried in the sand so that they were hidden from view while animals were foraging. Snail shells stuffed with pieces of shrimp were placed at one of two fixed locations approximately 70 cm from the burrow’s location. Foraging paths to and from the location of the food were video recorded from above (Fig. S1).

During trials, stomatopods made tortuous paths away from their burrows until they located the food in the arena. After animals found the food, they generally executed a well-directed homeward path to the burrow’s location. If the burrow was not encountered at the end of the homeward path, a search behavior was initiated (Fig. 2B and Video S1). To differentiate homeward paths from ongoing arena exploration, paths from food locations were considered to be homeward paths when they did not deviate more than 90° from their initial trajectories for at least one-third of the beeline distance (the length of the straightest path) from the food location to the burrow. Search behaviors were determined to be initiated when animals turned more than 90° from their initial homeward path trajectories.

**Figure 2.**
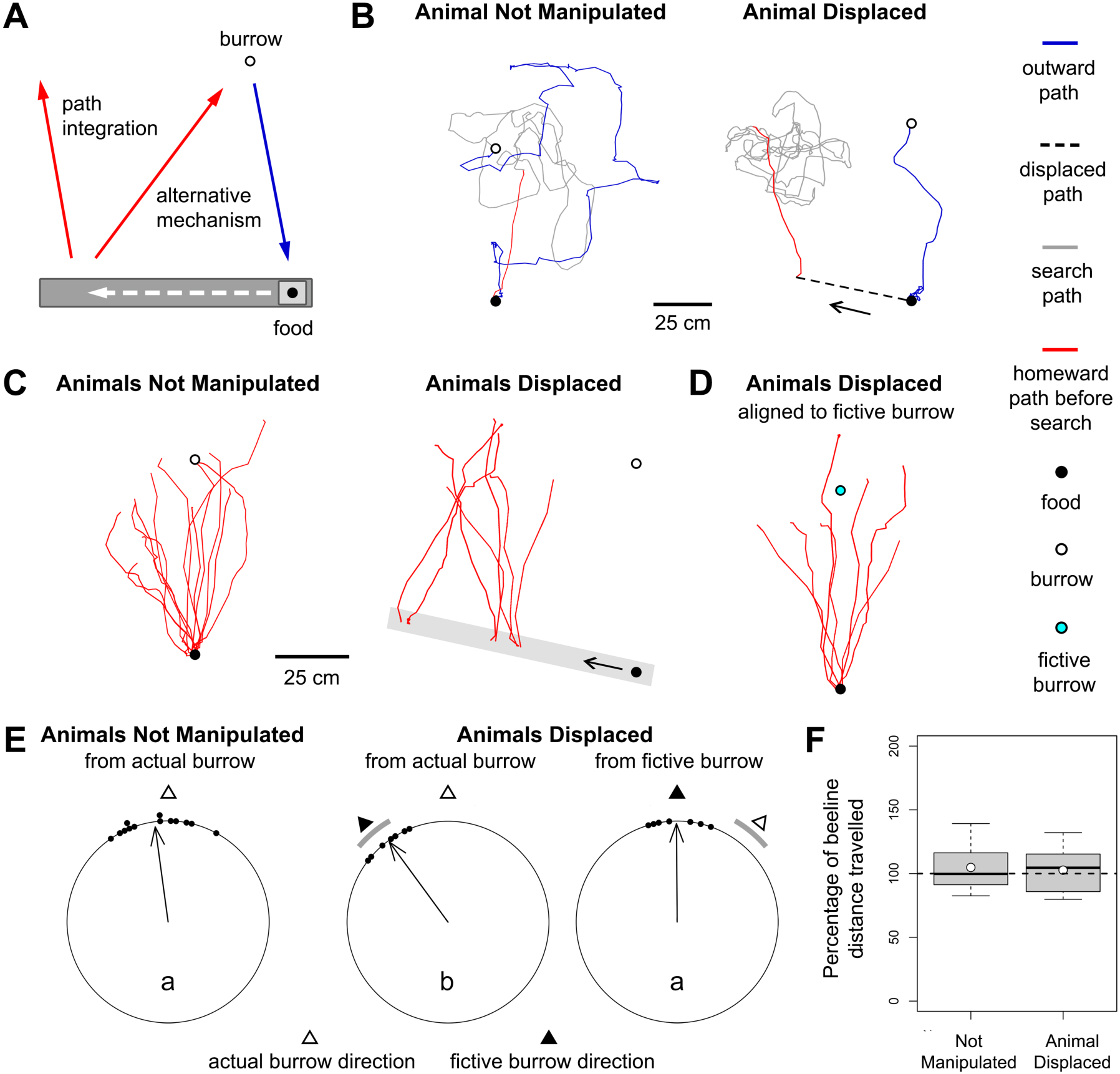
*Neogonodactylus oerstedii* uses path integration to navigate back to its burrow while foraging. (**A**) Experimental design. After passive displacement (dashed arrow), *N. oerstedii* should orient parallel to the direction of the burrow had it not been displaced if it uses path integration while homing. (**B**) Examples of foraging paths from and to the burrow when an animal was not manipulated and when an animal was passively displaced. (**C**) Data from all homeward paths. The grey rectangle represents the track on which animals were displaced and the empty circle represents the average burrow location with respect to the track. (**D**) Homeward paths when animals were displaced aligned to the position of the burrow had the animals not been displaced (the fictive burrow). (**E**) Orientations of homeward paths at one-third the beeline distance from the location of the food to the burrow. In all orientation diagrams, each point on the circumference of the circular plot represents the orientation of the homeward path of one individual with respect to either the actual position or the fictive position of the burrow. Arrows in each plot represent mean vectors, where arrow angles represent vector angles and arrow lengths represents the strength of orientation 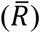. Different letters within orientation plots denote a significant difference between groups (p < 0.05). (**F**) The percentage of the beeline distance traveled during homeward paths before initiation of search behaviors. Data are from all trials in both conditions. Bars represent medians, points represent means, boxes indicate lower and upper quartiles, and whiskers show sample minima and maxima. The horizontal dashed line marks the beeline distance from the location of the food to the burrow.

Homeward paths of these animals were significantly oriented towards the burrow (−7.8° ± 5.36° (mean from the burrow where the burrow is 0° ± S.E.M.), P < 0.001; all statistical outcomes are presented in Tables S1 and S2; Fig. 2C, E). This observation suggested that *N. oerstedii* may use path integration to locate its burrow while forging.

To determine conclusively if *N. oerstedii* homes using path integration, homeward paths of foraging animals were observed after they had been passively translocated in the arena. This was accomplished by placing food on a thin platform on a track. Once an animal found the food, the platform (and animal) were carefully displaced along the track. If *N. oerstedii* path-integrate when foraging, animals should orient parallel to the direction of the burrow had they not been displaced (i.e. towards the fictive burrow location). Conversely, if animals oriented towards the burrow’s actual location, either *N. oerstedii* do not path-integrate but instead locate their burrows by some other method (perhaps using an odor emanating from the burrow or structural components of the greenhouse as orientation cues), or the experimental animals were aware of and integrated their passive displacement (Fig. 2A).

When animals were displaced, homeward paths were oriented towards the expected (fictive) location of the burrow rather than towards the actual location of the burrow (Fig. 2C, D and Video S2). Homeward orientations of stationary animals were significantly different from those of displaced animals measured in reference to actual burrow (p = 0.0016); however, they were not different when compared to homeward orientations of displaced animals measured in reference to the fictive burrow position (p > 0.1; Fig. 2E). Additionally, home vector lengths from both experiments were close to the beeline distance to the burrow (Not Manipulated: 104.9% ± 5.08% of beeline distance, Animal Displaced: 102.7% ± 7.3% of beeline distance (Mean ± Standard Error); Fig 2F). These results strongly support the hypothesis that foraging *N. oerstedii* use path integration when homing.

### Mantis shrimp orient using celestial and idiothetic cues during path integration

Path integration requires that an animal possess a compass to determine its headings and an odometer to measure the distances it travels. Due to the abundance of potential cues available for orientation, we examined whether the compass of *N. oerstedii* relies on cues external (allothetic) or internal (idiothetic) to the body. To distinguish between these two potential classes of cues, a rotatable platform was centered in a 1.5-meter diameter circular arena placed outdoors in an open, level field. Food was placed on the center of this platform 60 cm from the burrow. Trials were conducted in three environmental conditions: under clear skies, under partly cloudy skies when the sun was hidden by clouds, and under heavily overcast skies when celestial cues were obscured (Fig. S3). Trials were video recorded from cameras on tripods from above.

Once an animal found the food, the platform was carefully rotated 180°. If *N. oerstedii* used an allothetic compass for orientation, homeward paths should be oriented towards the location of the burrow, despite the animals’ passive rotation. Alternatively, if an idiothetic compass was used, homeward paths after rotation would be oriented approximately in the opposite direction (Fig. 3A). During additional trials, the tripods placed over the arenas were rotated either approximately 60° or 180° when animals were not manipulated to control for their presence as a potential orientation cue.

**Figure 3.**
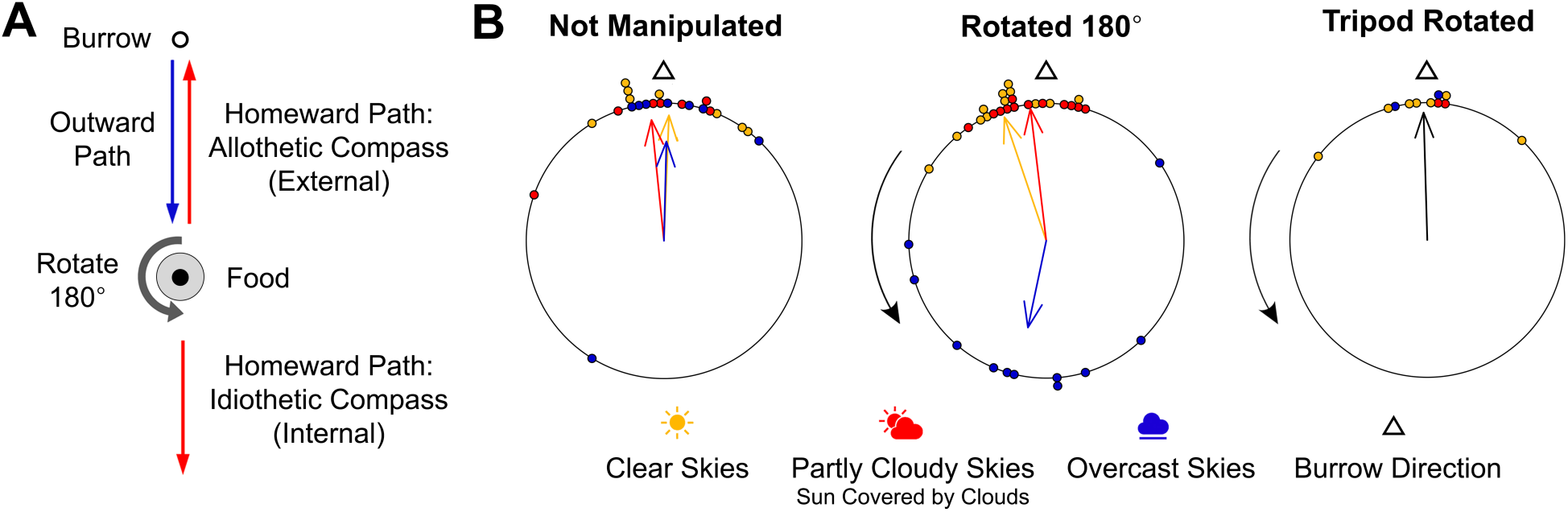
*Neogonodactylus oerstedii* uses celestial and idiothetic compasses during path integration. (**A**) Experimental design. Homeward paths using allothetic compasses were predicted to be oriented towards the burrow while homeward paths using idiothetic compasses were predicted to be oriented away from the burrow after passive 180° rotation. (**B**) Orientations of homeward paths when skies were either clear, partly cloudy with the sun covered by clouds, or heavily overcast. Animals were either not manipulated, rotated 180° on the platform, or the tripod was rotated during a trial (the direction of rotation is indicated by the black arrows). All groups exhibited significant orientations (p < 0.01).

When animals were not manipulated, they oriented significantly towards the burrow under all celestial conditions (clear: 3.27° ± 9.14°, P < 0.001, partly cloudy: −7.45° ± 11.54°, P < 0.001, overcast: −12.93° ± 20.27°, P = 0.001). During experiments in which animals were rotated on the central platform, animals continued to orient towards the burrow under clear (−18.66° ± 5.54°, P < 0.001) or partly cloudy skies (−5.88° ± 4.56°, P < 0.001). In contrast, when animals were rotated under heavily overcast skies, homeward paths were oriented away from the burrow (−168.02° ± 17.2°, P < 0.001). Finally, during trials when only the tripod was rotated, homeward paths were oriented towards the burrow (−1.48° ± 7.06°, P < 0.001; Fig. 3B and Videos S3-5).

These results indicate that when possible, *N. oerstedii* uses celestial cues for orientation; however, when celestial cues are obscured, *N. oerstedii* relies on an idiothetic compass. Under clear skies, *N. oerstedii* may use the solar azimuth for orientation. However, even when the sun was obscured, *N. oerstedii* continued to orient correctly despite being rotated as long as patches of clear sky were visible. Potential orientation cues used under this condition may include polarization patterns [5, 7, 13-16], spectral gradients [17, 18], or luminosity gradients in the sky [19, 20].

### Mantis shrimp use the sun as a compass while orienting

We determined if *N. oerstedii* uses the solar azimuth as a cue for orientation, using the outdoor arenas described above. Trials were conducted when the sun was clearly visible in an open sky at an altitude between 20° and 45° above the horizon. When animals had reached food placed in the center of the arena, the actual location of the sun was blocked by a board and the sun was instead reflected 180° from its original position in the sky by a mirror (Fig. 4A). If *N. oerstedii* uses a sun compass for orientation, under this condition homeward paths should be oriented away from the burrow (Fig. 4B). In control trials, the sun was concealed by the board but was not mirrored.

**Figure 4.**
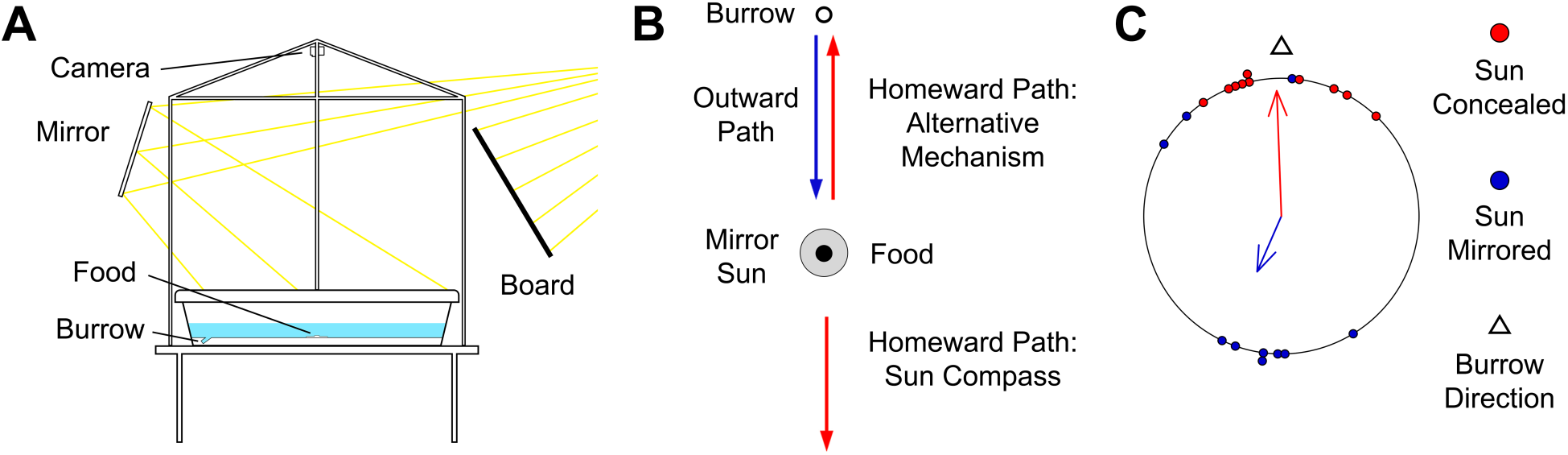
*Neogonodactylus oerstedii* uses the solar azimuth as a compass cue for orientation. (**A**) Sun compass arena design. (**B**) The sun was mirrored once animals reached the food location. Homeward paths were predicted to be oriented in the direction opposite to the burrow if *N. oerstedii* orients using the solar azimuth. (**C**) Orientations of the homeward paths when the sun was either concealed with a board or both, concealed with a board and mirrored to the opposite side of the arena. Both groups exhibited significant orientations (p < 0.05).

During experiments when the sun was concealed and mirrored 180° from its original location, homeward paths were primarily oriented in the opposite direction of the burrow (−156.3° ± 8.14°, P = 0.023, Video S6); however, during control trials when the sun was only concealed, homeward paths were oriented towards the burrow (−2.11° ± 4.27°, P < 0.001; Fig. 4C). These results indicate that *N. oerstedii* can use the solar azimuth for orientation. Nevertheless, even when the sun was mirrored, three of ten animals ignored the mirrored sun and correctly oriented home. These individuals either primarily orient using a cue other than the solar azimuth (in contrast to the majority of individuals tested) or some other factor may have caused other orientation cues to be ranked over the solar azimuth in these cases.

### Mantis shrimp orient using overhead polarization patterns

Since individuals were able to orient correctly after passive rotation under partly cloudy skies when the sun was obscured by clouds or when the sun was concealed by a board, *N. oerstedii* appears to use celestial cues other than the solar azimuth for orientation. Celestial polarization patterns, which are widely used by animals [5, 7, 13-16] and are clearly visible underwater at the depth ranges *N. oerstedii* occupies in nature [21], may have been used under these conditions. To determine if *N. oerstedii* can orient using overhead polarization patterns, indoor arenas were constructed over which an artificial polarization field was created using white LEDs and a composite filter constructed of a polarizer and a diffuser (Fig. 5A and Fig. S4). When the polarizer side of the composite sheet faced the arena, light passing through the filter created a linearly polarized light field. When the diffuser side faced downwards, a depolarized light field resulted (Fig. S4C-E). Thick black curtains were placed around the arena to create an isolated, nearly homogenous environment.

**Figure 5.**
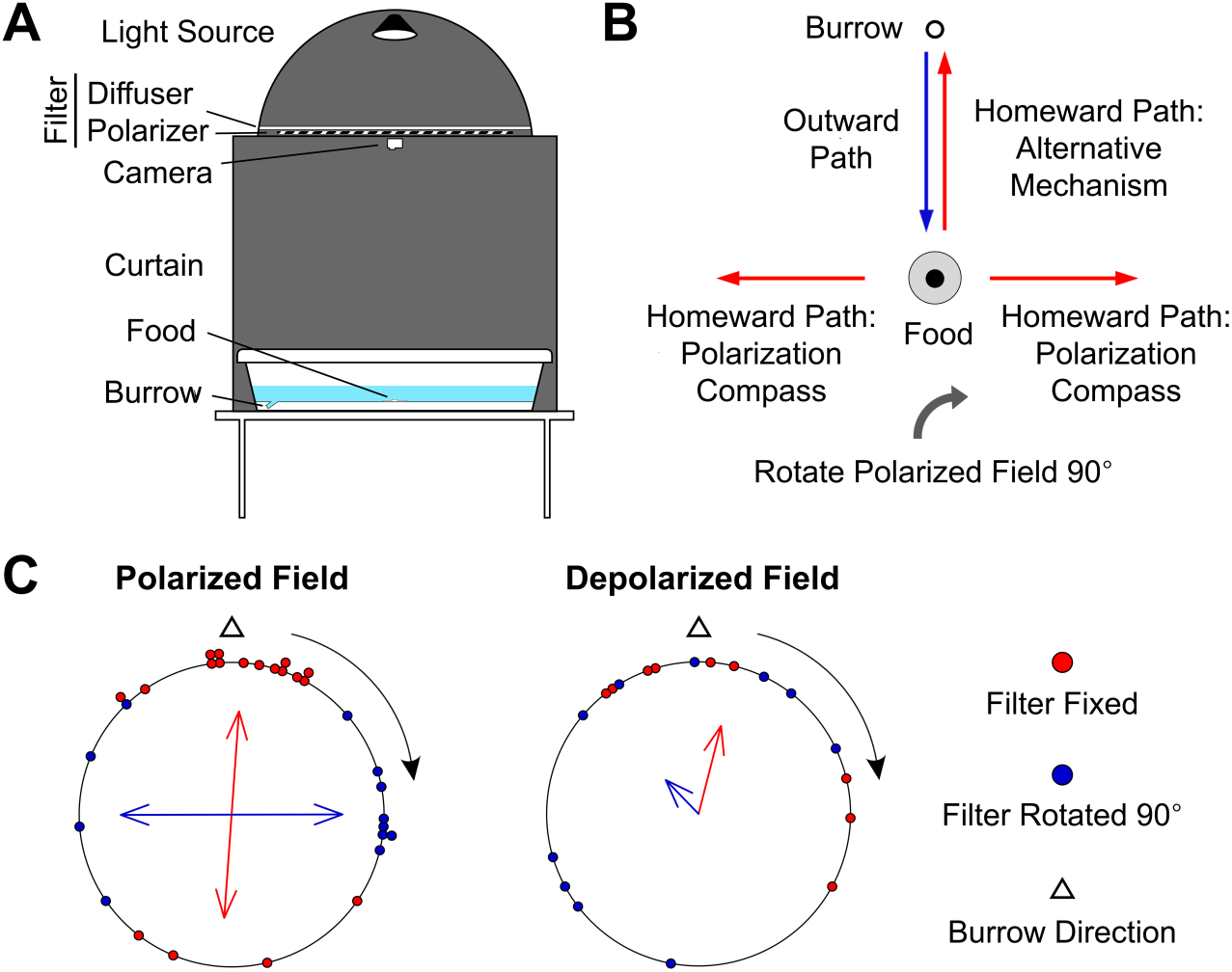
*Neogonodactylus oerstedii* uses overhead polarization patterns as a compass cue for orientation. (**A**) Polarization arena design. (**B**) The polarized light field was rotated 90° from its original position once animals reached the food location. Homeward paths were predicted to be oriented perpendicular to the direction of the burrow if *N. oerstedii* were using overhead polarization patterns for orientation. (**C**) Orientations of the homeward paths at one-third the beeline distance from the location of the food to the burrow when the overhead filter was oriented either with the polarizer or diffuser facing down towards the arena. Under each condition, the filter was either fixed in place or rotated 90° in the direction of rotation indicated by the black arrows. Doubled polarized field data can be reviewed in Extended Data Fig. 6. Both groups exhibited significant orientations under polarized fields (p < 0.01) and were either weakly or not significantly oriented under depolarized fields (fixed: p = 0.037, rotated: p = 0.39).

To manipulate the polarization field during experiments, once an animal found food centered in the arena, the polarized filter was rotated 90° from its original position. In this condition, if *N. oerstedii* used the overhead polarization field as a compass, homeward paths should be oriented perpendicular to the direction of the burrow (Fig. 5B). In control trials, the polarizer remained fixed throughout the experiment. During these experiments, homeward paths should be oriented towards or opposite to the burrow’s location (since a polarized light pattern is bidirectionally ambiguous). To control for the rotation of the filter, the filter was positioned to provide a depolarized light field and was rotated 90° when animals were at the location of the food. If experimental animals were only using the polarization field for orientation, homeward path orientations between the static and rotated depolarized light fields should not differ (Fig. S4B).

When the polarized field was static, homeward paths were oriented parallel to the direction of the burrow (data were doubled to create a unimodal distribution (see methods), 10.31° ± 34.03°, P < 0.001); however, when the polarized field was rotated 90°, individuals oriented their homeward paths perpendicular to the direction of the burrow (doubled data, 179.91° ± 13.74°, P < 0.001, and Video S7). Under a depolarized field, homeward paths were either weakly oriented or exhibited no significant orientation (static: 14.38° ± 19.59°, P = 0.037, rotated: −44.35° ± 25.35°, P = 0.39; Fig. 5D and Fig. S5). These results indicate that *N. oerstedii* uses overhead polarization patterns in the visible spectrum to orient.

In the outdoor experiments, animals oriented in the opposite direction to their burrows after being rotated 180° under heavily overcast skies, indicating that they relied on an idiothetic compass. Since animals were poorly oriented under indoor depolarized light fields, visual information may be important for idiothetic orientation in *N. oerstedii*. The photic environment in the indoor arenas differed mainly from the outdoor environment in two ways: UV light was absent in the indoor arenas, and the light available under overcast skies was over 50 times as bright as that available indoors (Fig. S4A-B). Visual information may be compromised indoors under these conditions. We also observed that home vector lengths were more varied under indoor conditions when compared to trials under natural lighting (P = 0.019, F = 2.75, indoor: n = 27, outdoor: n = 23; Fig. 6). This suggests that visual information such as rotational and translational optic flow fields might play a role in path integration in *N. oerstedii*.

**Figure 6.**
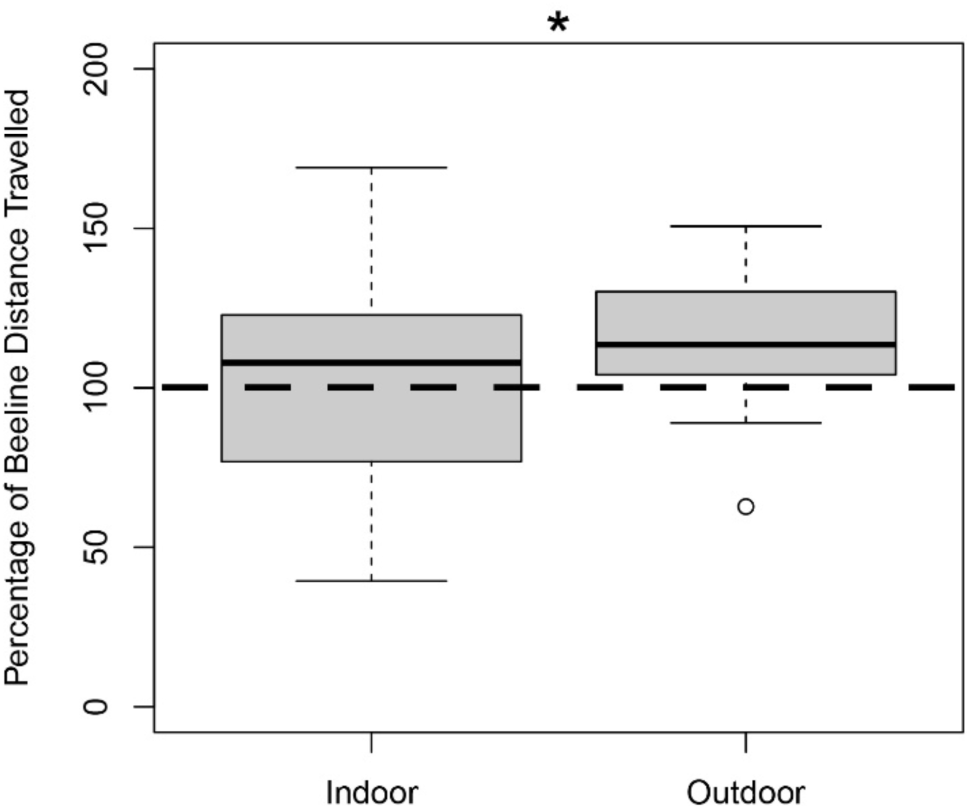
Lengths of homeward paths before search behaviors were initiated were more varied under indoor conditions than under outdoor conditions. The y-axis represents the percentage of the beeline distance from the food location to the burrow traveled during homeward paths before search behavior were initiated. The data included are from all trials that occurred outdoors under open skies and indoors in the polarization arenas when animals were not manipulated. The bar represent the median, the box indicates lower and upper quartiles, whiskers show sample minima and maxima, and the dot represents an outlier. The horizontal dashed line marks the beeline distance from the location of the food to the burrow.

### Hierarchy of compass cues

Our findings reveal a ranked hierarchy of cues used in the compass of *N. oerstedii*. During experiments in which the sun was mirrored, celestial polarization patterns were not affected, yet the majority of animals oriented relative to the displaced sun. This indicates that when the sun is visible in the sky, it is the primary compass cue used by *N. oerstedii*. However, when the sun is obscured, *N. oerstedii* orients using celestial polarization patterns (and potentially other celestial cues not tested). Finally, when no celestial cue is available, *N. oerstedii* appears to turn to idiothetic cues for orientation (Fig. 7). The use of idiothetic or other non-celestial cues for orientation may be particularly advantageous in the aquatic environment, where turbidity, wave action, and the attenuation of celestial cues due to absorption and scattering in the water column may disrupt visual orientation cues. Like many other animal navigators [15, 22-24], *N. oerstedii* uses multiple redundant cues for orientation to navigate successfully.

**Figure 7.**
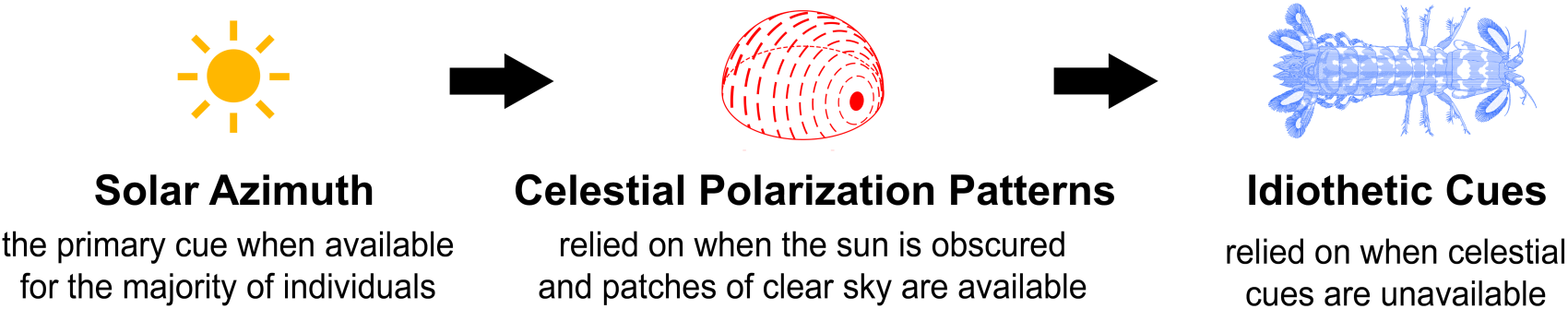
Proposed hierarchy of compass cues during path integration in *Neogonodactylus oerstedii*.

## Conclusion

Our results are the first to demonstrate path integration in a fully aquatic animal. Comparing the sensory cues involved in path integration in *Neogonodactylus* to those used by its terrestrial counterparts can provide insight into how navigational problems are solved in disparate environments with varying properties and challenges. Further, mantis shrimp occupy a wide variety of marine habitats, from clear tropical reefs to silty mud flats. Celestial cues easily viewed through the air-water interface in calm, shallow water are increasingly obscured with depth, turbidity, and wave action. While *N. oerstedii*, which mostly occupies shallow tropical waters, primarily relies on celestial cues for orientation, compass cue preferences likely differ for deeper-water stomatopod species or those that inhabit rougher, more turbid waters. The Earth’s magnetic field, a cue available throughout the water column, is known to be particularly useful for marine navigators [25-27] and may be used by stomatopods for orientation when celestial cues are unreliable as well.

This research opens a new avenue for the study of the neural basis of navigation in crustaceans, where insights into the evolution of arthropod brain structures and navigational strategies may be found. In insects, a highly conserved region of the brain called the central complex is thought to play a major role in navigation and path integration [28-32]. The neural organization of stomatopod central complexes is remarkably similar to those of insects [33]. Investigation of the function of neuropils within stomatopod central complexes could help uncover the evolutionary origins of navigation behaviors and the neural architecture of the central nervous system within arthropods; specifically the Pancrustacea, a taxon including both insects and malacostracan crustaceans, which is thought to have diverged from other arthropods over 600 million years ago [34].

## Acknowledgements

We thank N.S. Roberts, J. Park, S. Hulse, and V. Simmons for research assistance. We are grateful to J. Cohen and the University of Delaware, CEOE for space for experimentation.

## Author Contributions

R.N.P. designed and conducted all research, analyzed all data, and prepared the manuscript. T.W.C. provided guidance and research support.

## Data and Materials Availability

The data that support the findings of this study are available from the corresponding author upon reasonable request. Correspondence and requests for materials should be addressed to R.N.P.

## Materials and Methods

### Field Observations

*Neogonodactylus oerstedii* burrows were located in Florida Bay, offshore of the lower Florida Keys, USA. Burrows were video recorded from above using GoPro HERO 4 Black Edition cameras (GoPro Inc.) mounted to PVC tripods.

### Animal Care

Individual *Neogonodactylus oerstedii* collected in the Florida Keys, USA were shipped to the University of Maryland, Baltimore County (UMBC). Animals were housed individually in 30 ppt sea water at room temperature under a 12:12 light:dark cycle for indoor experiments and under local light:dark cycles (Lewes, DE, USA) for outdoor experiments. Animals were fed whiteleg shrimp, *Litopenaeus vannamei*, once per week.

Data were collected from 13 individuals during the initial experiments in the greenhouse and during translocation experiments (5 male and 8 female), 14 individuals during the outdoor rotation experiments (6 male and 8 female), 13 individuals during the sun compass experiments (7 male and 6 female), and 18 individuals during the zenithal polarization compass experiments (11 male and 7 female). All individuals were between 30 and 50 mm long from the rostrum to the tip of the telson.

### Experimental Apparatuses

#### Greenhouse Experiments

Four relatively featureless, circular navigation arenas were constructed from 1.5 m-diameter plastic wading pools that were filled with pool filter sand and artificial seawater (30 ppt, Fig. S1A). Arenas were placed in a glass-roofed greenhouse on the UMBC campus. The spectral transmittance of light through the greenhouse glass was nearly constant for all wavelengths, excluding the deep-UV-wavelength range (280 to 350 nm; Fig. S1B). Celestial polarization information was transmitted through the glass roof of the greenhouse (Fig. S1C). Vertical burrows created from 2 cm outer-diameter PVC pipes were buried in the sand 30 cm from the periphery of the arena so that they were hidden from view when experimental animals were foraging. Trials were recorded from above using C1 Security Cameras (Foscam Digital Technologies LLC) mounted to tripods placed above the arenas. During animal displacement experiments, a thin 11 x 82 cm acrylic track with a movable platform was placed 30 cm from the wall of the arena at its closest edge.

#### Outdoor Experiments

Outdoor navigation arenas were constructed from 1.5 m-diameter plastic wading pools with a white plastic base. 2 cm outer-diameter PVC pipe burrows were placed into holes drilled into the base of the arena 15 cm from the arena’s periphery. A rotatable platform was placed in the center of each navigation arena. Arenas were filled with filtered sea water (30 ppt). Trials were recorded from above using GoPro HERO 4 Black Edition cameras (GoPro Inc.) mounted to tripods placed above each arena. Arenas were placed in a wide empty open field at the University of Delaware’s College of Earth, Ocean, and Environment in Lewes, Delaware, USA.

During sun compass experiments, a rotatable 122 cm x 91 cm whiteboard on a vertical stand was used to block the sun while a 41 cm x 41cm glass mirror was used to reflect the sun to opposite side of the arena.

#### Indoor Experiments

Arenas used in the outdoor experiments were placed in a dark room. Arenas were surrounded by thick matte black curtains and lit from above using a centered diffused light source (Lepower 50W LED floodlights, 4000Lm, 6500K). Composite filters constructed of a linear polarizer (American Polarizers Inc., 38% transmission visible spectrum) and two sheets of wax-paper sandwiched between two sheets of colorless transparent acrylic were placed under each light source. When the polarizer side of the composite sheet faced downwards towards the arena, light was linearly polarized to an average degree of 99.91% from 420 to 700 nm. For unpolarized fields, the depolarizing waxed paper side faced downwards, reducing the average degree of polarization to 0.04% from 420 to 700 nm. The overhead polarization stimulus had an angular diameter of 27° of when viewed from the center of the arena.

### Experimental Procedures

Individual *N. oerstedii* were placed in each arena and were allowed to familiarize themselves to the arena for 24 hours. During familiarization, a vertical 2 cm diameter PVC column with alternating 1 cm thick black and white horizontal stripes was placed adjacent to the burrow, marking it during the animals’ initial explorations of the arena.

After familiarization, the column marking the burrow was removed from the arena. Empty *Margarites sp.* snail shells stuffed with pieces of food (whiteleg shrimp) were placed at fixed locations in the arena. During experiments conducted in the greenhouse, food was placed at one of two locations 50 cm from the periphery of the burrow. During experiments in which animals were displaced, food was placed on the movable platform on which animals were translocated. For the outdoor and indoor compass experiments, food was placed on rotatable platforms in the center of the arenas. Each animal was allowed three successful foraging excursions (i.e. food placed in the arena was found) before foraging paths were used for analyses. If an individual did not successfully locate food within one week in the arena, it was replaced with a new individual.

During animal translocation experiments, once animals found food placed on the movable platform, they were carefully displaced along the track to a new location in the arena by the pulling of a thin fishing line tethered to the platform.

During outdoor rotation experiments, trials were conducted under three environmental conditions: clear skies, partly cloudy skies when the sun was hidden by clouds, and heavily overcast skies. Full-sky polarization images were initially used to determine when trials would be categorized in the heavily overcast condition. It soon became clear that the lack of a visually identifiable solar disk during an overcast day (with clouds completely covering the sky) was sufficient to categorize a trial as occurring under a heavily overcast sky. Therefore, this method was used for the majority of celestial condition designations. During these experiments, animals were either not manipulated, carefully rotated 180°, or the tripods recording trials were rotated either approximately 60° or 180° to control for their presence. During experiments when animals were rotated 180°, animals were carefully rotated by the pulling of thin fishing line tethered to the platform once the animal found food placed on the rotatable platform. Animals were randomly rotated either clockwise or counterclockwise. During data analysis, clockwise trials were flipped so all trials could be analyzed in a counterclockwise fashion. During trials controlling for the presence of the tripod over the arena, the tripod was rotated when the animal was at the food location.

During outdoor solar azimuth compass experiments, trials were run only when the sun was clearly visible at an altitude between 20° and 45° above the horizon. When animals had reached food placed in the center of the arena, the location of the sun was blocked by a board and reflected 180° from its original position in the sky by a mirror. To control for the presence of the board, trials were also conducted when the sun was only blocked by the board but not mirrored.

During indoor experiments in which the polarization field was manipulated, once an experimental animal found food placed in the center of the arena, the composite polarization filter with the polarizer-side facing downwards over the arena, creating a linearly polarized field, was rotated 90° from its original position. In control trials, the polarizer was touched by the researcher, but remained fixed throughout the experiment. In order to control for the rotation of the filter, the filter was placed wax-paper-side face downward over the arena, creating an unpolarized field, and was rotated 90° when animals were at the location of the food.

### Data and Statistical Analyses

Foraging paths to and from food locations to the burrow were video recorded. In order to differentiate homeward paths from continued arena exploration, paths from the food locations were considered to be homeward paths when they did not deviate more than 90° from their initial trajectories for at least one-third of the beeline distance (the length of the straightest path) from the food location to the burrow. From these homeward paths, search behaviors were determined to be initiated when an animal turned more than 90° from its initial trajectory.

Paths were traced at a sampling interval of 0.2 seconds using the MTrackJ plugin [43] in ImageJ v1.49 (Broken Symmetry Software), from which the output is given as Cartesian coordinates. From these data, homeward path lengths and beeline distances from the food location to the burrow were determined. From these measures, home vector lengths were calculated as percentages of beeline distances.

Additionally, the orientation of homeward paths when animals were at one-third of the beeline distance from the food source to the burrow (at which point the orientation of the home vector was usually observed) was recorded using ImageJ.

All statistical analyses were run on R (v3.3.1, R Core Development Team 2016) with the “CircStats”, “circular”, “Hmisc”, and “boot” plugins. All statistical analyses for indoor polarization field experiments were performed subsequent to using the doubling angles procedure for bimodal data outlined in Batschelet (1981) [36]. Orientation data were analyzed using the following procedures for circular statistics [36].

All reported mean values for orientation data are circular means and circular standard errors of means.

Rayleigh tests of uniformity were used to determine if homeward paths were oriented within a group for the initial set of experiments and the translocation experiments run in the greenhouse. V-tests of uniformity were used to determine orientation in predicted directions of groups from the compass experiments (for outdoor rotation, sun compass, and zenithal polarization compass experiments).

Watson-Williams tests for homogeneity of means were used to determine if group orientations were significantly different from one another.

No significant difference was observed between homeward orientations of males and females during the initial set of experiments when animals were not manipulated (p > 0.5; Figure S2)) so data from both sexes were pooled for all experiments.

A F-test of equality of variances was used to determine if a difference in the variance of homeward vector path lengths could be observed between experiments run in the indoor arenas and experiments run outdoors when animals were not manipulated.

Bonferroni corrections were used for all tests when applicable. All statistical information including sample sizes, test statistics, P-values, means, and standard errors of means are presented in Tables S1 and S2.

### Polarization Imaging

A camera with a polaroid filter and a fisheye lens was used to take photographs of the sky near zenith for all full sky polarization images and zenithal polarization images of the indoor polarization arenas as per methods of Cronin et al. (2006) [37].

### Spectrometry

Irradiance measurements in the greenhouse were taken near midday on a cloudless day using an Ocean Optics USB2000 spectrometer connected to a 3 m long, 400 µm diameter, fiber-optic cable with a cosine-correcting head. The percentage of light transmitted through the greenhouse glass was calculated by comparing the ratio of irradiance measurements taken inside the greenhouse to irradiance measurements taken outside the greenhouse immediately afterwards. Irradiance measurements of light available in the indoor polarization arena and outdoors under heavily overcast skies were measured from the center of the arena using the same spectrometer system.

Transmittance measurements of the polarizing filter were taken using the same spectrometer system without a cosine-correcting head. The percent polarization of light transmitted through the filter was determined by calculating the ratio of the percent transmission of the polarizing filter layered with a second polarizer oriented first perpendicularly and then parallel to the first filter.

## Supplementary Information

**Figure S1.**
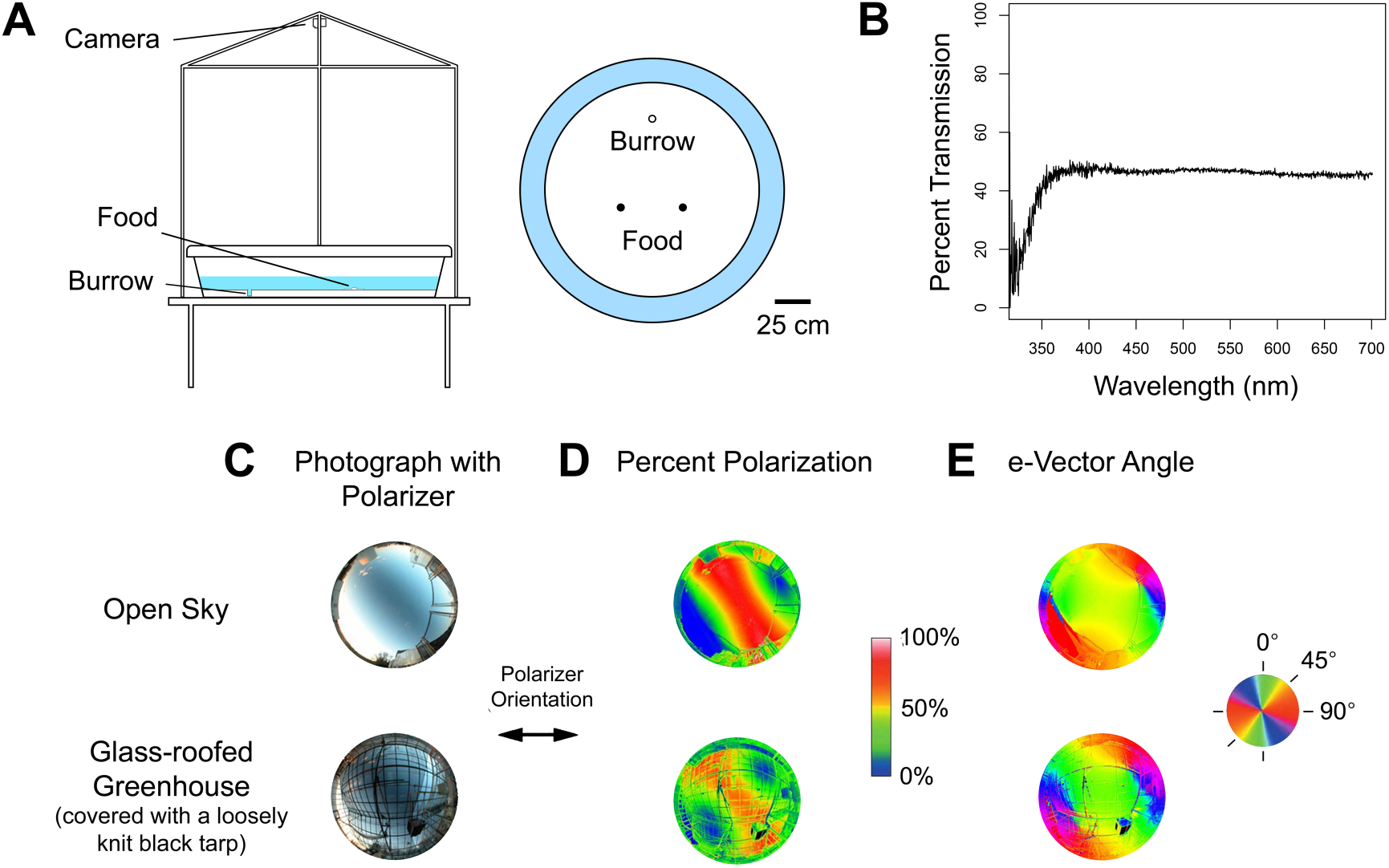
Greenhouse arena design. (**A**) Navigation arenas 150 cm in diameter contained a burrow (empty circle) buried in the base of the arena 30 cm from the arena’s periphery. During trials in the greenhouse when animals were not manipulated, food was placed at one of two positions 50 cm from the periphery of the arena (filled circles). Trials were video recorded from above. (**B**) Transmission of irradiance spectra through the glass-roof of the greenhouse on November 24, 2015 at 15:30. The spectral transmittance of light through the glass roof of the greenhouse is nearly constant for all wavelengths greater than ∼360 nm. (**C-E**) Celestial polarization patterns are transmitted through the glass roof of the greenhouse. (**C**) Photographs of the sky at sunset on a day with very few clouds (November 24, 2015) using a fisheye lens and linear polarizer set in the east-west direction (as indicated by the arrow in the legend). Photos were taken inside and outside the glass-roofed greenhouse used for the initial set of experiments. (**D**) Percent polarization. Warmer regions in the images indicate higher percent polarization and cooler regions indicate lower percent polarization (see key). (**E**) e-Vector angle, indicated by the color corresponding the key to the right of the images.

**Figure S2.**
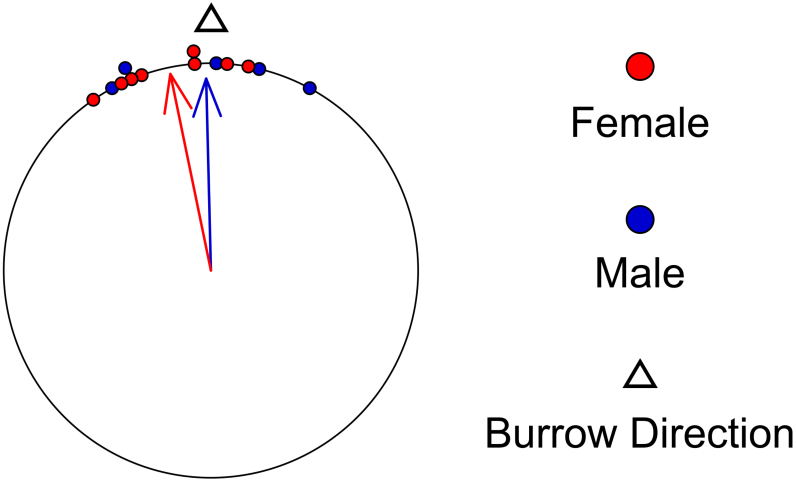
Male and female *N. oerstedii* orient towards home equally well while foraging. Homeward orientations of male and female individuals during experiments in the greenhouse when animals were not manipulated. Each point along the circumference of the circular plot represents the orientation of the homeward path of one individual with respect to position of the burrow (empty triangle). Blue-filled circles represent males while red-filled circles represent females. Arrows represent mean vectors, where angles of the arrows represent the mean vector angles and arrow lengths represent the strength of orientation in the mean direction 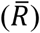. Males (n=5) and females (n=8) both exhibited significant orientations (p < 0.01 for both groups). No significant difference in orientation was observed between males and females (p>0.5).

**Figure S3.**
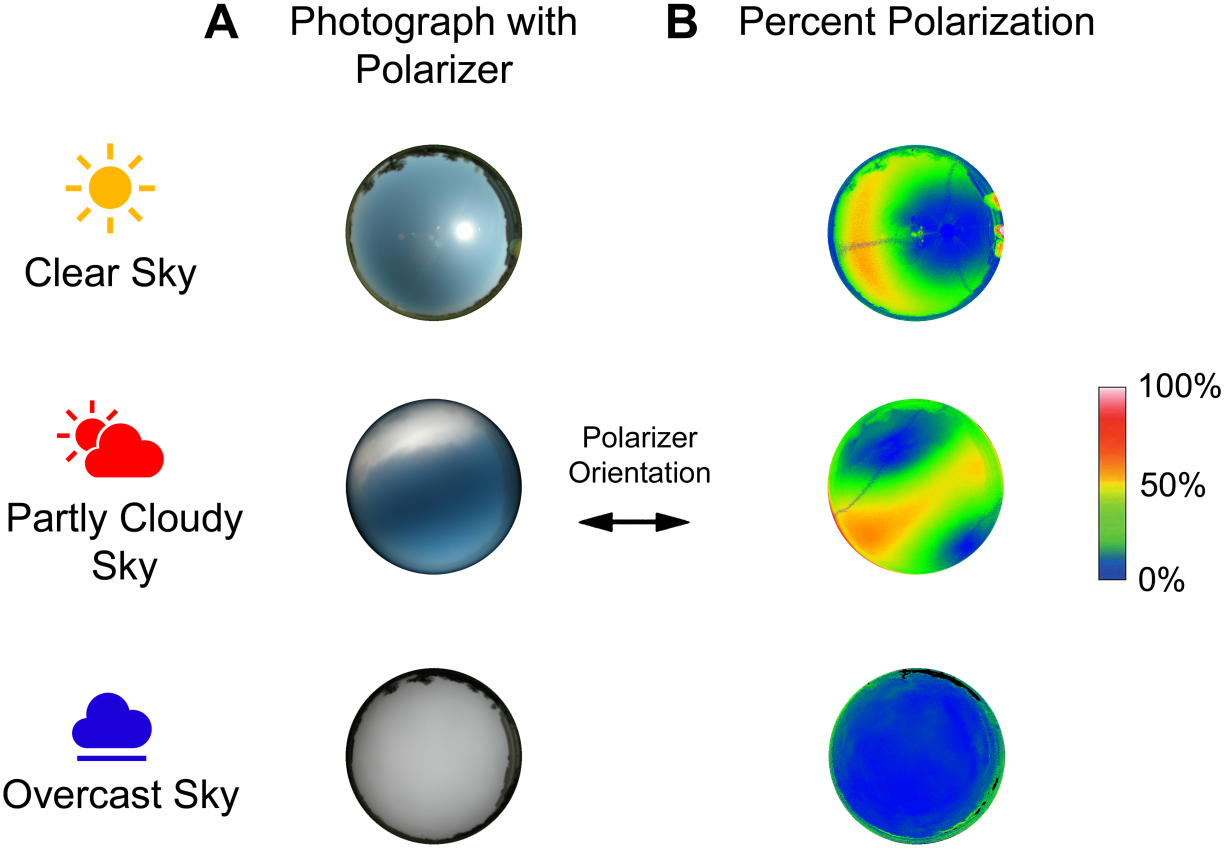
Celestial conditions during outdoor rotation experiments. (**A**) Photographs of the sky on days with clear, partly cloudy (when the sun is covered by clouds), and heavily overcast skies in June 2018 taken using a fisheye lens and linear polarizer set in the east-west direction (as indicated by the arrow in the legend). Photos were taken in a field at the University of Delaware’s College of Earth Ocean and Environment in Lewes, DE. (**B**) Percent polarization. Warmer regions of the images indicate higher percent polarization and cooler regions indicate lower percent polarization (see key).

**Figure S4.**
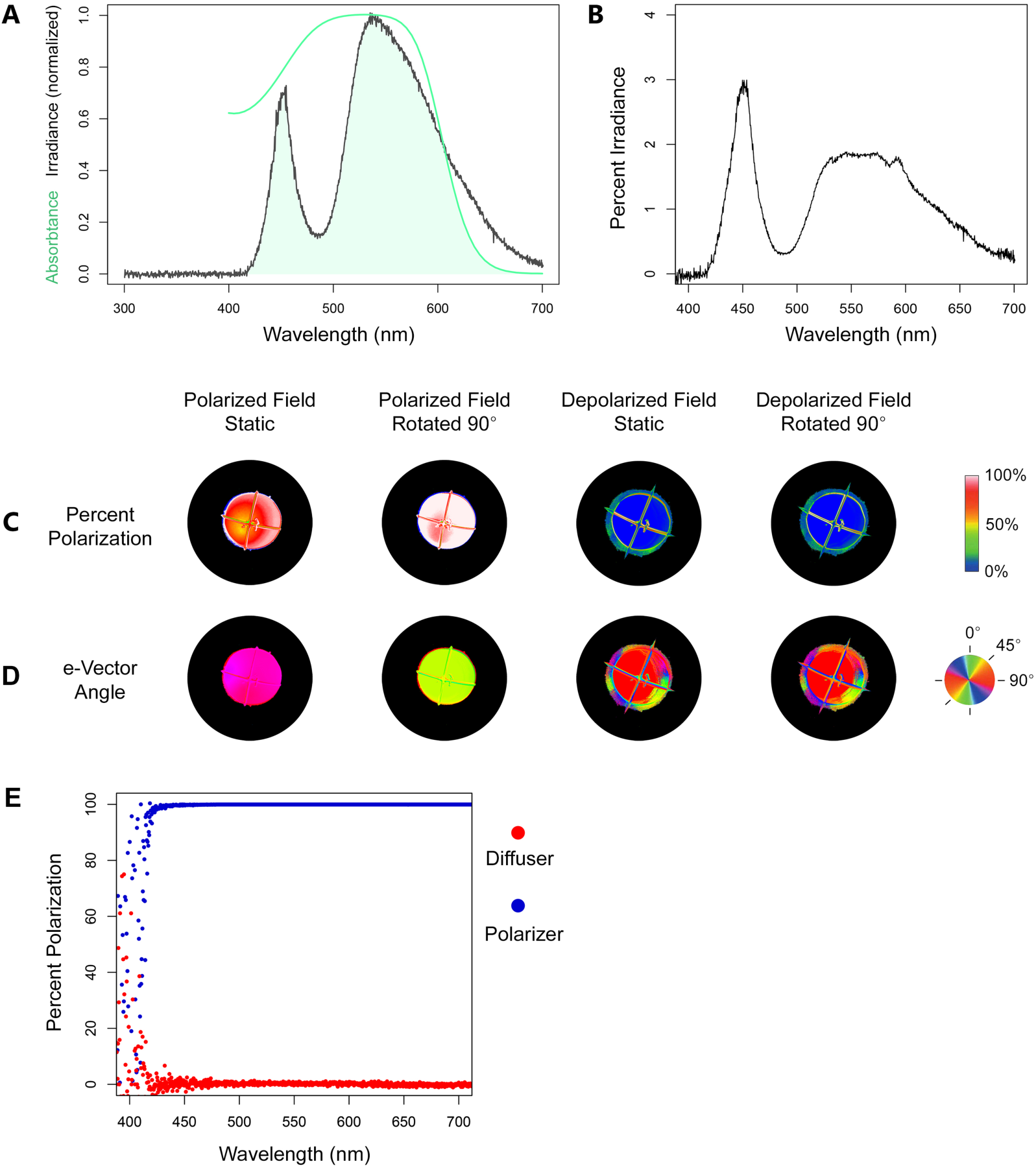
Photic conditions in the indoor polarization arenas. (**A**) Irradiance spectra of light available in the indoor polarization arenas (black line). Absorptance curve of the main rhabdomeric photoreceptors (R1-7) in the peripheral hemispheres of the eyes of *Neogonodactylus oerstedii* (green line; from Cronin and Marshall, 1989 [38]). The shaded area represents light available in the arena absorbed the main rhabdoms in the peripheral hemispheres of the eye of *N. oerstedii*. (**B**) Percent irradiance of the indoor polarization arenas compared to an overcast sky. The light environment during outdoor experiments under overcast skies was over 50 times as bright as during those run in the indoor arenas at 540 nm (the brightest wavelength in indoor arenas). **(C-E)** Polarization information of the overhead feature for all experimental conditions of the indoor polarization experiments. (**C**) Percent polarization of an indoor artificial polarization field arena near the zenith. Warmer regions of the images indicate higher percent polarization and cooler regions of the images indicate lower percent polarization (see key). (**D**) e-Vector angles of the same images as in a. e-Vector angle is indicated by the color corresponding to the key on the right of the images. (**E**) Percentage of polarized light transmitted through the overhead filter used in the indoor polarization experiments when either the polarizer side (blue) or diffuser side (red) faced down over the arena.

**Figure S5.**
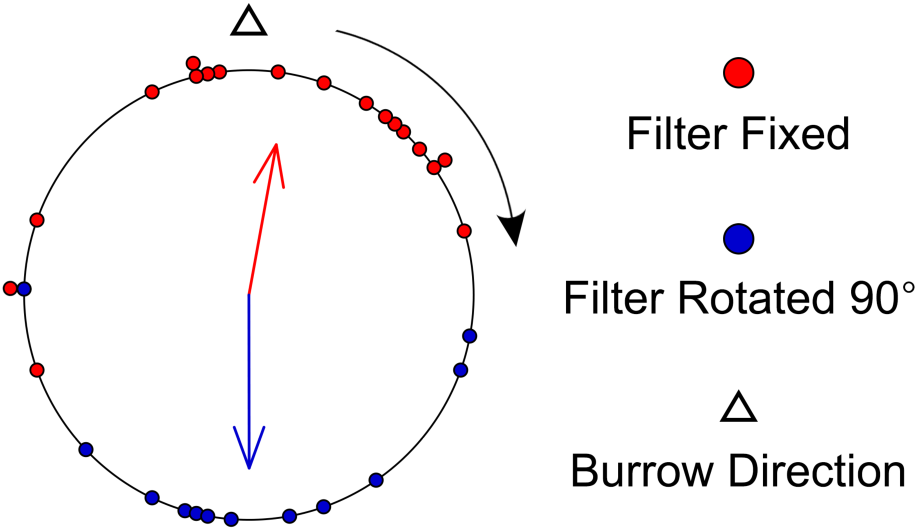
Doubled data for the polarized light field experiments. Orientations of homeward paths under a polarized field at one-third the beeline distance from the location of the food to the burrow after the data was doubled to create unimodal distributions (see methods). Red-filled points represent trails when the overhead filter was fixed in place while blue-filled points represent trials when the filter was rotated 90° with the direction of rotation indicated by the black arrow. Each point along the circumference of the circular plot represents the orientation of the homeward path of one individual in respect to the position of the burrow (empty triangle). Arrows in each plot represent mean vectors, where direction of the arrow represents the mean vector angle and arrow length represents the strength of orientation in the mean direction 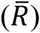.

**Table S1:**
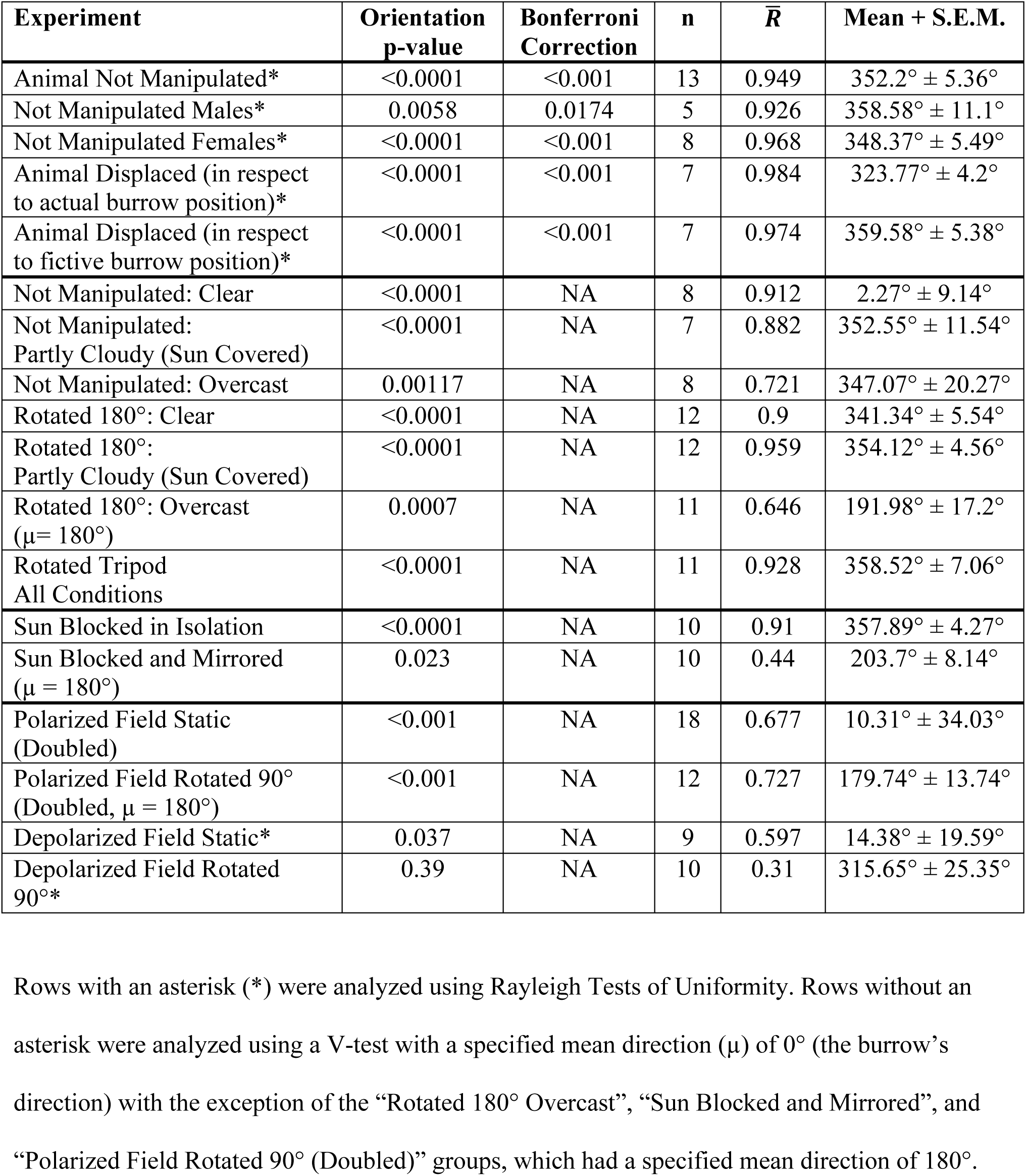
Summary of orientation statistics for all experimental groups.

**Table S2:** Summary of Watson-Wheeler Tests of Homogeneity of Means.

| Experiment | p-value | Bonferroni Correction | F |
| --- | --- | --- | --- |
| Not Manipulated: Males vs. Females | 0.3767 | 1.0 | 0.8487 |
| Not Manipulated vs. Animal Displaced (in respect to actual burrow position) | 0.00016 | 0.00048 | 13.783 |
| Not Manipulated vs. Animal Displaced (in respect to fictive burrow position) | 0.383 | 1.0 | 0.8 |

**Video S1. Foraging behavior of *Neogonodactylus oerstedii* showing homing resulting from path integration.** Outward path is in blue, home vector path is in red, and search path is in grey. Filmed at 30 frames per second. Replay speed is indicated in the bottom-right corner of the video.

**Video S2. *Neogonodactylus oerstedii* orients parallel to the direction of home after it has been displaced.** Outward path is in blue, displaced path is in yellow, home vector path is in red, and search path is in grey. Filmed at 30 frames per second. Replay speed is indicated in the bottom-right corner of the video.

**Video S3. *Neogonodactylus oerstedii* orients towards home after being rotated 180° under a clear sky.** Outward path is in blue, rotated path is in yellow, and home vector path is in red. Filmed at 30 frames per second. Replay speed is indicated in the bottom-right corner of the video.

**Video S4. *Neogonodactylus oerstedii* orients towards home after being rotated 180° under a partly cloudy sky.** Outward path is in blue, rotated path is in yellow, and home vector path is in red. Filmed at 30 frames per second. Replay speed is indicated in the bottom-right corner of the video.

**Video S5. *Neogonodactylus oerstedii* orients away from home after being rotated 180° under a heavily overcast sky.** Outward path is in blue, rotated path is in yellow, and home vector path is in red. Filmed at 30 frames per second. Replay speed is indicated in the bottom-right corner of the video.

**Video S6. *Neogonodactylus oerstedii* orients away from home after the sun was concealed by a board and mirrored to other side of the arena.** Outward path is in blue and home vector path is in red. Filmed at 30 frames per second. Replay speed is indicated in the bottom-right corner of the video.

**Video S7. *Neogonodactylus oerstedii* orients perpendicular to the direction of home after a polarized field over the arena was rotated 90°.** Outward path is in blue and home vector path is in red. Filmed at 30 frames per second. Replay speed is indicated in the bottom-right corner of the video. Apparent patterns in the center of the arena are reflections of the light source on the water’s surface.

